# A Novel DNA-binding Protein Coordinates Asymmetric Chromosome Replication and Chromosome Partitioning

**DOI:** 10.1101/091496

**Authors:** James A. Taylor, Gaël Panis, Patrick H. Viollier, Gregory T. Marczynski

## Abstract

Bacterial chromosome replication is regulated from a single replication origin (*ori*) that receives cell cycle signals. Following replication, bacteria often use the *parABS* partition system with a centromere-like *parS* locus to place the chromosomes into the daughter cells. Our knowledge of cell cycle regulation is incomplete and we searched for novel regulators of chromosome replication. Here we show that in the cell cycle model *Caulobacter crescentus* a novel DNA-binding protein promotes both the initiation of chromosome replication and the earliest step of chromosome partitioning. We used biochemical fractionation to identify a protein (OpaA) that preferentially binds to mutated *ori* DNA that also increases *ori*-plasmid replication *in vivo*. OpaA represents a previously unknown class of DNA-binding proteins. *opaA* gene expression is essential and sufficient OpaA levels are required for the correct timing of chromosome replication. Whole genome ChIP-seq identified the genomic binding sites for OpaA, with the strongest associations at the *parABS* locus near *ori*. Using molecular-genetic and fluorescence microscopy experiments, we showed that OpaA also promotes the first step of chromosome partitioning, the initial separation of the duplicated *parS* loci following *ori* replication. This separation occurs before the *parABS* mechanism and it coincides with the regulatory step that splits the symmetry of the chromosomes so that they are placed at distinct cell-poles which develop into replicating and non-replicating cell-types. We propose that OpaA coordinates replication with the poorly understood mechanism of early chromosome separation. *opaA* lethal suppressor and antibiotic experiments argue that future studies be focused on the mechanistic roles for transcription and translation at this critical step of the cell cycle.

**Author Summary:** Like all organisms, bacteria must replicate their chromosomes and move them into the newly dividing cells. Eukaryotes use non-overlapping phases, first for chromosome replication (S-phase) followed by mitosis (M-phase) when the completely duplicated chromosomes are separated. However, bacteria combine both phases so chromosome replication and chromosome separation (termed chromosome “partitioning”) overlap. In many bacteria, including *Caulobacter crescentus*, chromosome replication initiates from a single replication origin (*ori*) and the first duplicated regions of the chromosome immediately begin “partitioning” towards the cell poles long before the whole chromosome has finished replication. This partitioning movement uses the centromere-like DNA called *“parS”* that is located near the *ori*. Here we identify a completely novel type of DNA-binding protein called OpaA and we show that it acts at both *ori* and *par*S. The timing and coordination of overlapping chromosome replication and partitioning phases is a special regulatory problem for bacteria. We further demonstrate that OpaA is selectively required for the initiation of chromosome replication at *ori* and likewise that OpaA is selectively required for the initial partitioning of *par*S. Therefore, we propose that OpaA is a novel regulator that coordinates chromosome replication with the poorly understood mechanism of early chromosome separation.

## Introduction

Faithful genome duplication requires dedicated mechanisms to coordinate cell cycle events but for bacteria such coordination is especially problematic. For example, bacteria often experience very high growth rates that force the evolution of cell cycle programs where chromosome replication and chromosome partitioning overlap [1]. Otherwise, their sequential phasing would take too long and severely reduce competitive fitness. Bacteria also navigate constantly shifting and often extremely variable environments. This navigation likewise places extreme demands on regulatory mechanisms, and especially those mechanisms that choose the critical time to initiate chromosome replication. Once started, the cell is committed to chromosome replication that lasts most of the cell division cycle. Replication timing is critical since delays also reduce competitive fitness while commitment in unfavorable conditions risks genome damage and cell death [2]. Since a bacterial chromosome uses only one origin of replication (*ori*), the bacterial strategy for regulating chromosome replication implies that much information is processed through an *ori* and this also implies many new protein-binding interactions [3, 4]. Our working hypothesis is that *ori* information processing may require novel DNA-binding proteins that evolved to coordinate multiple inputs on cell status and proliferation.

*Caulobacter crescentus* evolved in challenging aquatic environments and it provides an excellent model to study the bacterial cell cycle [5]. Its dimorphic growth presents distinct programs of chromosome replication and chromosome partitioning that are being exploited for detailed analysis. *C. crescentus* swarmer (*Sw*) cells present the motile and non-replicating cell stage. As the cell cycle proceeds, the *Sw* cells differentiate into stalked (*St*) cells that present the non-91 motile but replicating stage. Therefore, the initiation of chromosome replication is coordinated with the cell differentiation from the *Sw* to the *St* cell stages. Next, *C. crescentus* cell division proceeds asymmetrically as the elongating cell builds a new flagellum at the new *Sw* pole opposite the old *St* cell pole. Chromosome partitioning starts very soon after the initiation of chromosome replication and both cell cycle processes overlap the elaboration of asymmetric cell division that ultimately yields a *Sw* cell and a *St* cell [6]. Therefore, this cell division program produces distinct cells, each with distinct non-replicating (*Sw*) and replicating (*St*) chromosomes.

We study the *C. crescentus* chromosome origin of replication (*Cori*; Fig1A) and our experiments address how *Cori* directs the non-replicating *Sw* and the replicating *St* chromosome states. In this report, we identify a novel DNA-binding protein (OpaA) and show that it selectively facilitates the initiation of chromosome replication and the earliest stage of chromosome partitioning. Therefore, OpaA activity connects the start of two overlapping yet otherwise mechanistically different cell cycle processes that ultimately create two asymmetric chromosomes.

**Fig 1.**
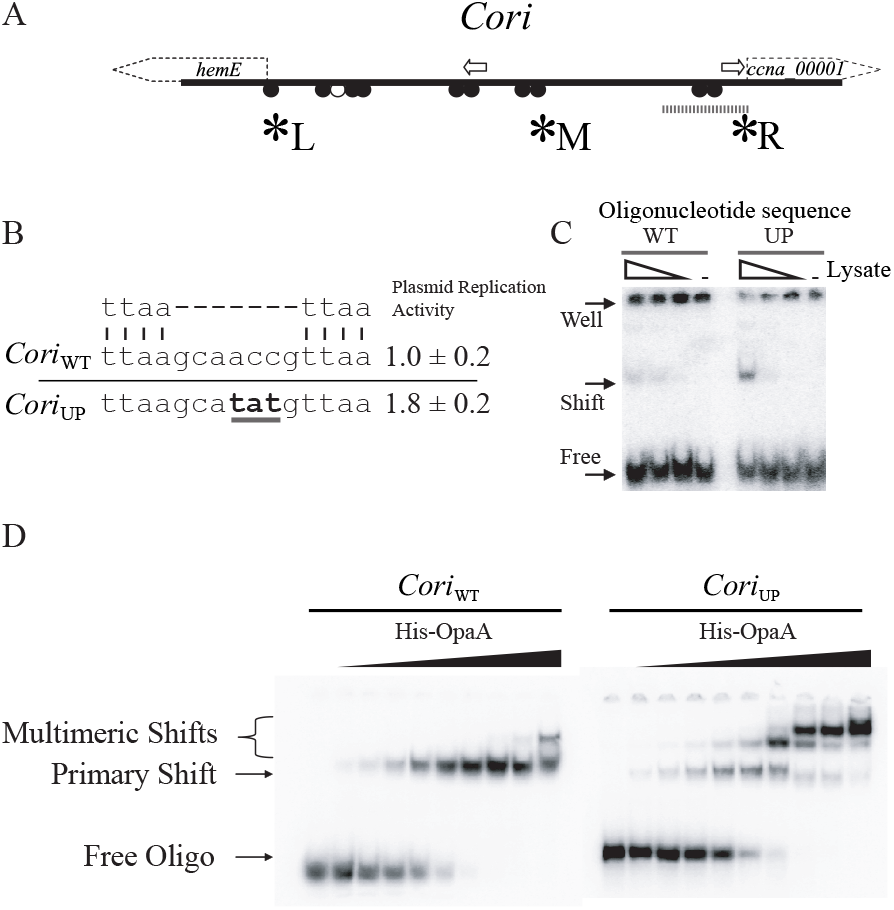
Identification of OpaA (CCNA_03428) as a factor promoting chromosome replication. A) A schematic of *Cori* including CtrA binding sites (black circles indicate half-sites), strong DnaA binding sites (two open arrows) and the most conserved region, studied in Taylor et al. 2011 (underline). Flanking coding sequences (*hemE* and *ccna_00001*) are shown. The positions of three probes used for qChIP analysis in Fig 3B are marked by asterisks together with their identifying letters, for *Cori* L, *Cori* M and *Cori* R. B) The ability of WT and mutant *Cori* to drive autonomous plasmid replication was assessed by activity of the *gusA* gene carried by the plasmid reporting plasmid copy number. The alignment shows WT sequence compared to up-replication mutant sequence (*Cori*UP; mutation underlined) together with the plasmid copy number (relative to WT). The position of a consensus CtrA binding site is indicated above the alignment. C) A factor present in a cleared *C. crescentus* cell lysate binds more strongly to oligonucleotides carrying sequences from *Cori*UP than from *Cori*WT. 40 bp annealed oligonucleotides carrying either the *Cori*WT or *Cori*UP sequences were incubated with ten-fold dilutions of partially fractionated cleared cell lysate (left to right, dilutions indicated by a wedge) or in buffer alone (lanes marked “-”) and run on a non-denaturing gel. A greater fraction of the *Cori*UP oligonucleotides are shifted by the same lysate concentrations than the *Cori*WT oligonucleotides. D) EMSA reactions in which equal concentrations of His-OpaA (from *E. coli*) were used to shift annealed oligonucleotides as in (C). For each gel, in the leftmost lane oligonucleotides were incubated in buffer alone. Reactions in the right hand lane were incubated with 500 nM His-OpaA, and lanes to the left of this contain a two-fold dilution series of His-OpaA.

## Results

### Discovery of a novel origin binding protein

As part of a search for new regulators of the *C. crescentus* origin of chromosome replication (*Cori*) we previously reported a 5 bp mutation within the most conserved region of *Cori* (Fig 1A) that causes a ~2 fold increase in *Cori* autonomous replication [7]. Here, we introduced a 3 bp mutation into a similar position in *Cori* and observed that this mutation also caused a ~2-fold increase in *Cori* replication (Fig 1B). We hypothesized that this mutation (*Cori*_UP_) altered the binding of a protein to *Cori*, and we sought to identify it by an electrophoretic mobility shift assay (EMSA). We probed *C. crescentus* cell lysates using radio-labeled annealed oligonucleotides carrying either the natural *Cori* sequence (*Cori*_WT_) or the mutated up-replication sequence (*Cori*_UP_). We found that the *Cori*_UP_ sequence was shifted with higher affinity than the *Cori*_WT_ sequence by a protein in the lysate (Fig 1C). We purified this protein by biochemical fractionation (see S1 Fig and methods) and identified it using liquid chromatography/tandem mass-spectrometry as the product of the *ccna_03428* gene, previously annotated as a hypothetical cytosolic protein [8], which we re-named as the origin and partition locus gene (*opaA*). We PCR cloned and purified N-terminally poly-histidine tagged OpaA from *E. coli* and confirmed that this protein preparation also differentially shifts the *Cori*WT and *Cori*UP 123
oligonucleotides (Fig 1D).

### OpaA is essential in *C. crescentus*

We attempted to delete the *opaA* coding sequence and to replace it with the omega antibiotic resistance cassette through two-step homologous recombination. However, we could only delete *opaA* when we supplied an extra *opaA* gene on a replicating plasmid (see S1 Table). A high-throughput transposon mutagenesis screen also proposed that *opaA* is essential in *C. crescentus* [8]. To study how removing *opaA* might affect *C. crescentus*, we replaced the plasmid covering the deletion with a plasmid carrying *opaA* under the control of a xylose inducible promoter (pJT157; see S2A Fig). This strain (GM3817) now required xylose for growth (S2B Fig) and allowed us to deplete OpaA by shifting from media containing xylose to media containing glucose.

### OpaA is required for timing chromosome replication

Our first analysis indicated that OpaA positively promotes replication by directly binding to *Cori* (Fig 1), and therefore we next tested replication initiation in conditionally expressing *Pxyl::opaA* GM3817 cells. Chromosome replication initiates from *Cori* when swarmer (*Sw*) cells differentiate into stalked (*St*) cells. Therefore, synchronous *C. crescentus Sw* cell cultures of WT parental GM1609 and GM3817 were isolated [9] and placed in media supplemented with xylose or glucose. Samples of the synchronized cultures were removed at 0, 30, 60 and 90 minutes post-synchronization and treated with antibiotics to prevent new rounds of chromosome replication and cell division. Since this treatment allows ongoing chromosome replication to finish, this method distinguishes replicating and non-replicating cells. Cells that initiated chromosome replication when sampled will complete replication (but not cell division), and will have two complete chromosomes and cells which have not initiated replication will retain only one chromosome (see Fig 2A).

**Fig 2.**
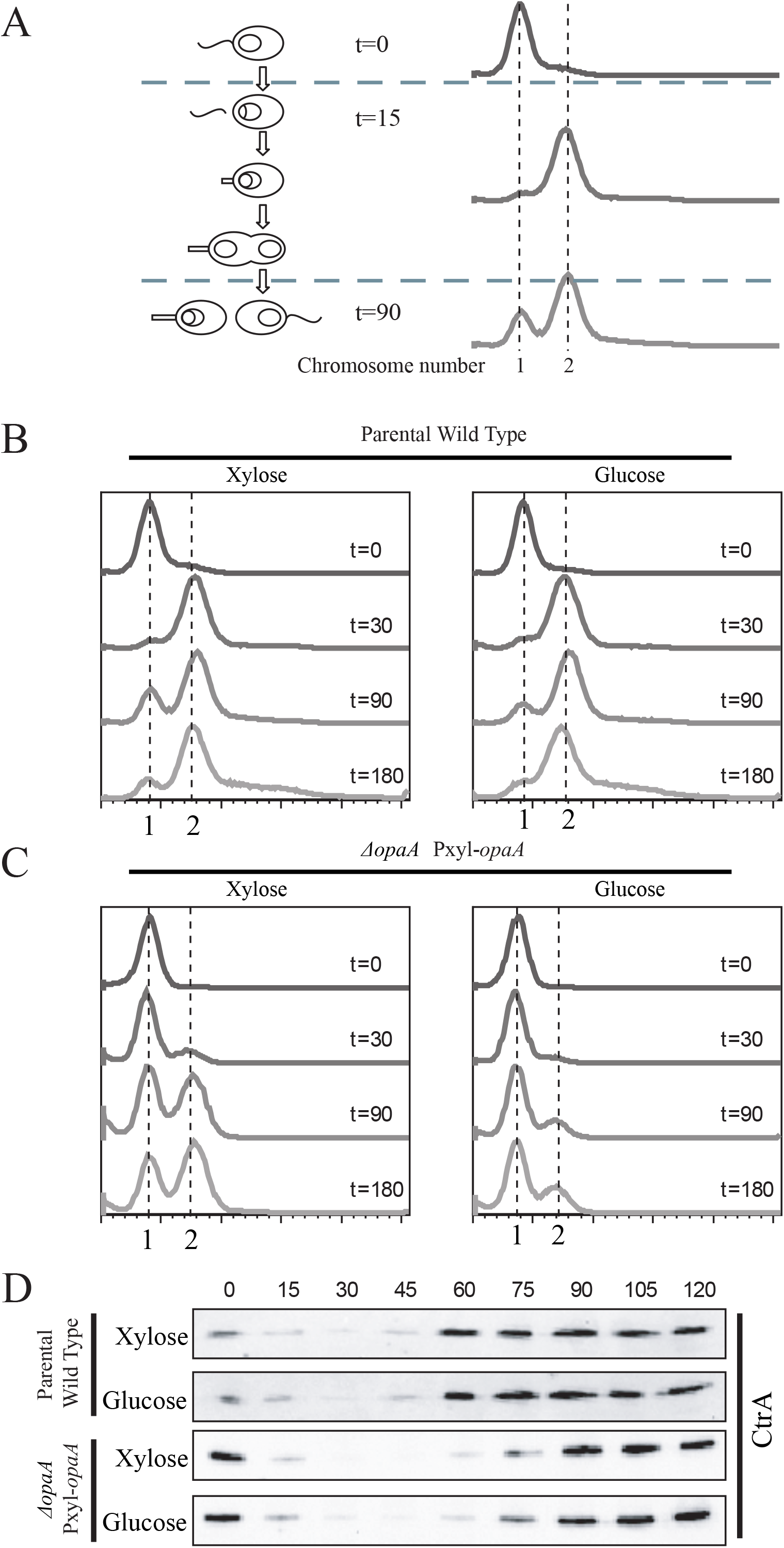
Cells with insufficient OpaA fail to initiate chromosome replication. A) Presentation of the flow cytometry-based assay to measure initiation times of chromosome replication. A synchronized culture (of wild type strain GM1609) proceeds through the cell cycle (cartoon on left) developing from swarmer cells (top) to stalked cells, predivisional cells, and finally divided cells (bottom). Flow cytometry is used to assess the replication state of the chromosome. Cells are transferred at various time points into media containing antibiotics that block further replication initiation or cell division events and are allowed to finish rounds of chromosome replication that have started (see *Methods*). Flow cytometry can then be used to assess the replication state of the chromosome (traces on right). Prior to initiation, all cells contain one chromosome, while after initiation all cells contain two chromosomes. B and C) Initiation time in synchronized cultures of the parental wild type strain (GM1609; B) and the OpaA Pxyl conditional-expression strain (GM3817; C) assessed using flow cytometry as outlined in (A) grown in media containing xylose (left histograms; OpaA expression induced) or glucose (right histograms; OpaA expression not induced). Samples were taken at the indicated times (minutes). D) Western blots for CtrA master regulator protein with samples taken at time points indicated above the lanes (minutes) from synchronized cultures in (B) and (C) indicate that their cell cycles, with respect to cell differentiation, are otherwise proceeding normally.

As expected, cells from WT GM1609 differentiated rapidly and initiated chromosome replication by the 30 minute time point and GM1609 showed no replication timing differences between xylose or glucose media (Fig 2B). In contrast, even with xylose to induce OpaA, GM3817 cells displayed a pronounced delay in replication initiation, and most of these cells failed to initiate replication in glucose media without xylose to induce OpaA expression (Fig 2C). Western blots show that OpaA levels in GM3817 grown in xylose were ~10% of the WT cells (not shown). Therefore, both replication delays (WT versus GM3817 Fig 2BC and xylose versus glucose Fig 2C) are attributable to lower OpaA levels.

We also assayed the synchronized cells in Fig 2 by tracking the abundance of 2 marker proteins that are degraded at the *Sw* to *St* cell transition. Western blot analysis demonstrated that both the master cell cycle regulator CtrA (Fig 2D) and the chemotaxis protein McpA (Fig S3) were degraded at the appropriate cell cycle times. For example, CtrA was completely degraded in all cultures by 30 min when all WT GM1609 cells had initiated chromosome replication. Interestingly, both marker proteins show a delayed return late in the cell cycle of the GM3817 cells suggesting that later stalked cell programs (e.g. chromosome partitioning and cell division) are delayed with reduced OpaA levels. In summary, while OpaA probably aids later cell cycle progression, it does not significantly influence the earlier *Sw* to *St* cell differentiation and yet lower OpaA levels selectively retard the timely start of chromosome replication, implying the selective regulation of *Cori.*

### Direct OpaA binding to *Cori* promotes chromosome replication *in vivo*

If OpaA is promoting chromosome replication by directly binding to *Cori*, then increasing *Cori* affinity for OpaA should ameliorate the delayed replication seen above during OpaA depletion. We therefore created the strain GM3880 with the *Cori*UP mutation placed at the natural *Cori* position but which is otherwise identical to GM3817 used above in Fig 2. By using GM3880 in the otherwise same replication timing experiment, we now see that the *Cori*UP mutation advances replication initiation towards that of the WT GM1609 control and the severe replication timing lag is no longer seen in glucose media (compare flow cytometry results in Fig with S4A Fig).

Similarly, if increased OpaA binding to *Cori* is driving the extra *Cori*-plasmid replication of *Cori*UP seen in Fig 1B, then genetically removing OpaA should remove the differential replication of the *Cori*WT and CoriUP sequences. To test this prediction, we isolated a spontaneous suppressor strain which retained viability despite lacking *opaA* (GM3875; see S1 Text, methods). Repeating the autonomous replication assays confirms that the ~2-fold differential replication between the *Cori*WT and *Cori*UP *Cori*-plasmids seen in WT GM1609 is no longer seen in GM3875 (S4B Fig).

### Genomic binding of OpaA also suggests functions at the chromosome partitioning locus

We used chromatin-immuno-precipitation and Illumina sequencing (ChIP-seq) to identify how OpaA is distributed on the *C. crescentus* chromosome in growing WT GM1609 cells. To show that ChIP-seq specifically identifies OpaA binding positions, we performed the same analysis on the spontaneous suppressor mutant GM3875. This comparison confirmed the specificity of our OpaA anti-serum, because none of the major binding peaks of WT GM1609 are present in the *opaA* GM3875 control (Fig 3A; compare outer black trace with inner grey Δ trace). Interestingly, we found that the most OpaA DNA-binding occurs around the chromosome partition locus (Fig 3A and Table 1), which is located within 10 kb of *Cori*. The next two most abundant peaks are placed between *Cori* and the partition locus *parABS* and at *Cori* itself. We independently confirmed abundant OpaA binding at these key positions using ChIP and qPCR (qChIP; S5A Fig). In addition to its role in replication initiation, these results suggest that OpaA also has a role(s) in chromosome partitioning, which we investigate further below.

**Fig 3.**
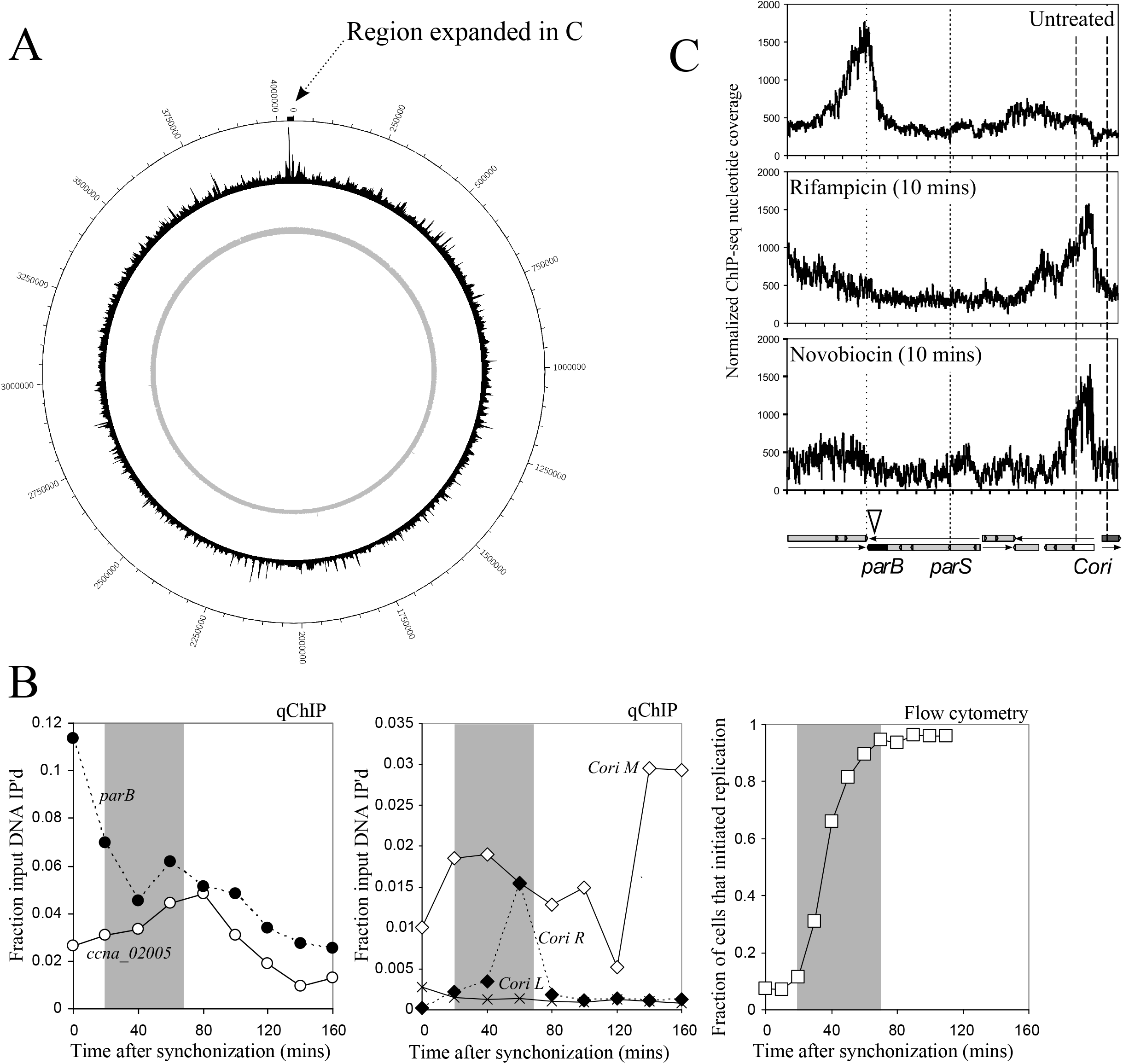
Global binding pattern of OpaA. A) The circular *C. crescentus* chromosome aligned two whole-genome ChIP seq signals (175 bp resolution histogram bins representing % of total reads) using the OpaA anti-serum first on the WT strain (GM1609, black outer signal) and then the *opaA* null suppressor strain (GM3875, grey inner signal). The standard bp positions of the WT NA1000 genome are displayed in the outer circle, with the small black bar above the 0-position indicating the region that is expanded in (C). B) qChIP (OpaA anti-serum) analysis performed on synchronized WT GM1609 cells grown in M2G media. OpaA binding was measured by qPCR at the following positions: at *parB* (left panel; filled circles, dashed line), at *ccna_02005* (left panel; open circles, solid line), and at three positions in *Cori* (middle panel, see Fig 1A for positions of probes: Cori L, crosses, solid line; Cori M, open diamonds, solid line; Cori R, filled diamonds, dashed line). The grey window shows the cell cycle period during which 95% of the chromosomes initiated replication, as measured by flow cytometry (right panel; see Fig 2A and *Methods*). C) ChIP-seq signals over the region surrounding *parB* and *Cori* from untreated WT GM1609 cells (top trace) and from the same culture sampled and treated for 10 minutes with 30 ug/ml rifampicin (middle trace), or 100 ug/ml novobiocin (bottom trace). In the schematic below these 3 ChIP seq signals, the genetic organization in this region is shown with the *parB* gene indicated in black, and *hemE* and *ccna_00001* which flank *Cori* indicated in white and dark grey respectively. The *Cori* region (the *Bam*HI fragment that defines the autonomously replicating *Cori*) is indicated by vertical dashed lines (long dashes), the position of *parS* by one vertical dashed line (short dashes) and the position of the intergenic region downstream of *parB* by a vertical dotted line. The white vertical arrowhead indicates the position of the *parB* primers used for the qChIP analysis in (B). Note that (A) and (C) represent data from whole-genome ChIP-seq experiments on independent cultures and that both experiments position all major whole-genome OpaA peaks at the same locations.

**Table 1.** The four most significant peaks in the OpaA genome ChIP-seq dataset.

| Rank | Start<sup>a</sup> | End<sup>a</sup> | Overlapping Genes | % Significant Reads<sup>*</sup> |
| --- | --- | --- | --- | --- |
| 1 | 4027601 | 4031150 | *leuS-holA-parB<sup>b</sup>* | 49.8 |
| 2 | 4038301 | 4040650 | *rho-ccna_03877* | 9.4 |
| 3 | 4040751 | 4042700 | *Cori(hemE-hemH)<sup>c</sup>* | 6.7 |
| 4 | 2152551 | 2154100 | *ccna_02005-02006* | 4.8 |

<sup>a</sup> – The NA1000 genomic coordinates of each peak. <sup>\*</sup> – The percentage of all the sequencing reads over the entire genome above the mean plus two standard deviations (see *Methods*) present in each peak. <sup>b</sup> – Convergent *leuS/holA* and *parAB* operons. <sup>c</sup> – Overlaps *Cori* replication sequences.

### DNA-binding properties of OpaA

OpaA probably binds DNA through very specific peptide contacts, because an OpaA mutation (Y82A) at a conserved tyrosine abolishes all high affinity EMSA binding to *Cori* (S6 Fig). We attempted to identify DNA sequences recognized by OpaA using the MEME software suite [10]. While this sequence alignment approach has identified DNA binding sites with similar ChIP-seq data [11], we did not find DNA motifs that accounted for the OpaA distributions on the chromosome. However, we noted that OpaA binding peaks often coincided with head-to-head transcription (Fig 3C and S7 Fig). We therefore tested whether transcription affects OpaA DNA-binding. Comparing ChIP-seq signals from untreated cultures and from cultures treated with rifampicin or novobiocin, we saw that most OpaA binding peaks were eliminated after 10 min with both treatments, as for example the peak at *parB* (Fig 3C). Only one major peak was not reduced: the peak at the left side of *Cori*, within the *hemE* gene, was increased after both antibiotic treatments. We used qChIP experiments with the same inhibitors to confirm this result. While ChIP-seq reports relative amounts of binding, our qChIP experiments demonstrate that absolute OpaA binding to *Cori* at *hemE* actually increases by approximately 50% when transcription is blocked, suggesting that OpaA is redistributed upon RNA-polymerase and upon DNA-gyrase inhibition (S5B Fig) and that special OpaA interacting DNA sequences lie at *Cori*. Although transcription affects OpaA binding to DNA, OpaA does not significantly affect RNA polymerase activity because the RNA levels of genes under the major OpaA binding peaks (Table 1) were not changed from WT GM1609 levels in the OpaA depletion strain GM3187 and in the *opaA* null suppressor strain GM3875 (data not shown). Therefore it is unlikely that OpaA regulates transcription by initiation or termination mechanisms. This negative result further suggests that OpaA acts selectively on chromosome replication and partitioning.

### OpaA binding correlates with the initiation of chromosome replication and partitioning

We examined OpaA binding during the cell cycle with qChIP experiments on synchronized WT GM1609 cells (Fig 3B). Regarding the 1st ranked ChIP-seq OpaA binding region (Table 1), with qChIP primers at *parB* this signal peaked early in the cell cycle immediately preceding the *Sw* to *St* cell transition and the initiation window for chromosome replication (20-60 min Fig 3B). This timing contrasts with the 4th ranked OpaA binding region, at *ccna_02005* close to the terminus, which peaked much later at ~80 min in to the cell cycle. We also examined OpaA binding at three positions spanning *Cori* (Fig 1A; asterisks). *Cori* L binding proximal to *hemE* was weak and showed no cell cycle regulation. In contrast, at *Cori* M binding to the middle of *Cori* was much stronger and this signal doubled during the initiation window for replication and again late in the cell cycle prior to the next round of replication initiation that starts after cell division. In contrast to these patterns, OpaA binding to *Cori* R, close to the initial binding site that we identified in Fig 1, peaked most discretely at 60 min, during the latter half of the initiation window for replication (Fig 3B). Therefore, OpaA binding changes significantly during the cell cycle: OpaA binding inside *Cori* correlates with the initiation of chromosome replication and the distinct *Cori* L M R binding patterns during replication initiation suggest specialized roles. However, peak OpaA binding at *parB* precedes chromosome duplication and separation and yet remains high, suggesting early roles before and during chromosome partitioning.

### OpaA is dynamically localized during chromosome partitioning

*C. crescentus* chromosome partitioning has been studied in *gfp-parB* strains where the GFP-ParB fusion protein binds the *parS* (centromere-like) locus near *Cori* to form fluorescent foci at the cell poles [12]. We fused the mCherry protein to the N-terminus of OpaA and integrated this construct at P*xyl* in strain GM3905 (*gfp-parB*) to create strain GM3921 (*gfp-parB Pxyl::mCherry-opaA*). This mCherry-OpaA protein is induced by xylose and it retains important *in vivo* functions, as GM3921 derived strains remained viable when the native *opaA* gene was deleted (see S1 Table). Therefore mCherry-OpaA localization probably reports the native OpaA localization. When we first examined an unsynchronized GM3921 culture we observed that mCherry-OpaA fluorescence was distributed as gradients (clouds) that changed with the apparent stage of the cell cycle (not shown). We therefore synchronized GM3921 cells and observed how the mCherry-OpaA distribution changed during the cell cycle (Fig 4 and S8 Fig). In t=0 *Sw* cells, mCherry-OpaA initially distributed as a gradient with its peak very near but not overlapping the flagellated cell pole marked by the single GFP-ParB focus. As chromosome replication and partitioning initiated, in t=15 and t=30 *St* cells, one GFP-focus remained polar while one GFP-focus moved away and the mCherry-OpaA peak moved with this partitioning GFP-ParB focus (peaks of GFP and mCherry fluorescence co-localized in 76% of partitioning cells at t=15). This movement created a wider mCherry-OpaA free gap between the polar GFP-focus and the partitioning GFP-ParB focus (seen in 94% of partitioning cells at t=15). The GFP-ParB focus then passed through the peak of the mCherry-OpaA gradient as it moved further away to the opposite pole. Finally in t=90 pre-divisional cells, mCherry-OpaA established a separate gradient in each nascent cell compartment, with a peak of mCherry-OpaA close to, but separated from, 275 each pole marked by its GFP-ParB focus.

**Fig 4.**
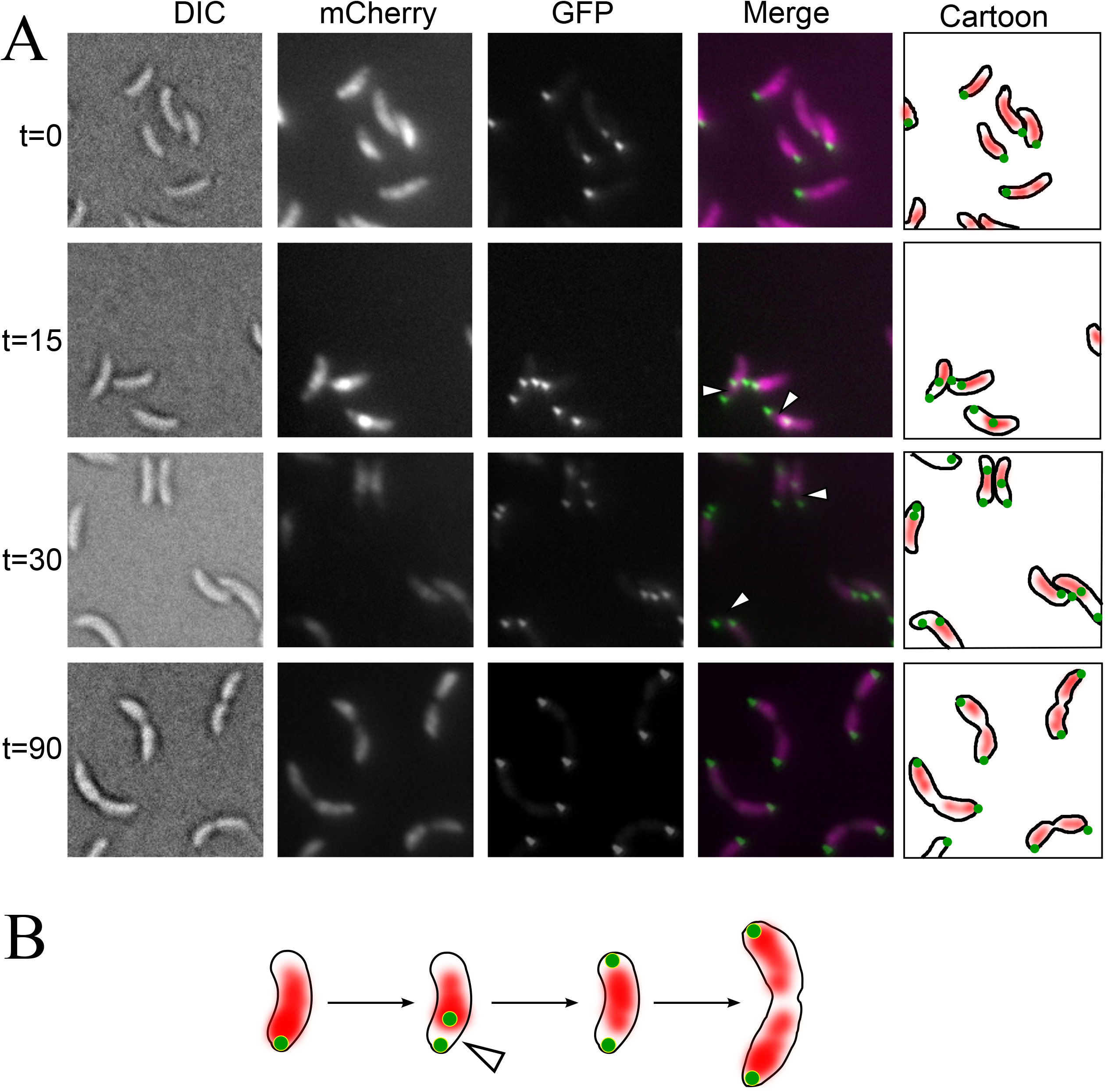
OpaA distribution changes during cell cycle progression and correlates with chromosome *parS* movements. A) Microscopy images from a synchronized culture of GM3921 (*gfp-parB Pxyl::mCherry-opaA*) taken at t=0, t=15, t=30 and t=90 (minutes). The full set of time points taken every 15 min is presented in S8 Fig. At each time-point we imaged >400 cells. Representative fields viewed with DIC, mCherry and GFP channels are shown in black and white and the merged mCherry/GFP channels are colored (pink and green respectively). The arrowheads in the merged t=15 and t=30 pictures mark cell regions between the two GFP-ParB foci that (compared with t=0 cells) become exceptionally free of mCherry-OpaA. In these cells the peak mCherry-OpaA fluorescence often co-localizes with the partitioning GFP-ParB focus. Cartoons of adjacent images show GFP-ParB (green) and mCherry-OpaA (red). B) A cartoon summary of inferred GFP-ParB and mCherry-OpaA movements over the cell cycle from *Sw* cell (t=0, left) to pre-divisional cell (t=90, right). As in (A) the arrowhead marks the sub-cell region between two GFP-ParB foci that becomes cleared of mCherry-OpaA as both fluorescence signals move together at this early phase of the cell cycle.

### Early-phase partitioning is assisted by OpaA

The dynamic cell cycle localization of mCherry-OpaA (Fig 4B), the early cell cycle binding (Fig 3B) as well as the 1st ranked magnitude (Fig 3A, Table 1) of OpaA binding to the partition region near *Cori* all suggest a role in chromosome partitioning. To help define this role, we performed OpaA depletion experiments in strains with chromosomal *gfp-parB* that were otherwise identical to those used above to study chromosome replication. Specifically, we compared GM3905 *gfp-parB* and GM3918 *gfp-parB ΔopaA Pxyl::opaA* and we observed abnormally localized GFP-ParB foci predominantly in GM3918 upon shifting glucose media (not shown). However, these GM3918 cells also showed cell division defects and since chromosome replication and partition are overlapping processes, we could not conclude that misplaced GFP-ParB foci directly resulted from OpaA depletion.

To search for consequences directly caused by OpaA, we took advantage of the δ*opaA* suppressor strains. Most importantly, we can select suppressor strains with normal replication kinetics during synchronous *Sw* to *St* differentiation (S4C Fig) and this allowed us to observe how OpaA and its absence could independently affect chromosome partitioning. Therefore, we compared synchronous cultures of a *gfp-parB ΔopaA* suppressor strain (GM3920) with the *gfp-parB* parental strain (GM3905) and we quantified 3 types of GFP-ParB localization patterns (unipolar, partitioning and bi-polar patterns, Fig 5ABC) at progressive cell cycle times. While bipolar GFP-ParB foci appeared at the same rate in both strains (Fig 5C), GM3920 showed more unipolar (Fig 5A) and fewer partitioning foci (Fig 5B) during the early 0 to 40 min cell cycle period. We therefore focused on this early partitioning period; we repeated these synchronies in triplicate and examined the distribution of GFP-ParB foci at the 20 min time point. This fixed time analysis (Fig 5D) confirmed the time-course analysis (Fig 5ABC) and demonstrated that GM3920 cells without OpaA have a specific partitioning defect, revealed by a deficit of double-foci cells during the early phase of partitioning, an intermediate phase between the unipolar (unpartitioned) and the final bi-polar positioning.

**Fig 5.**
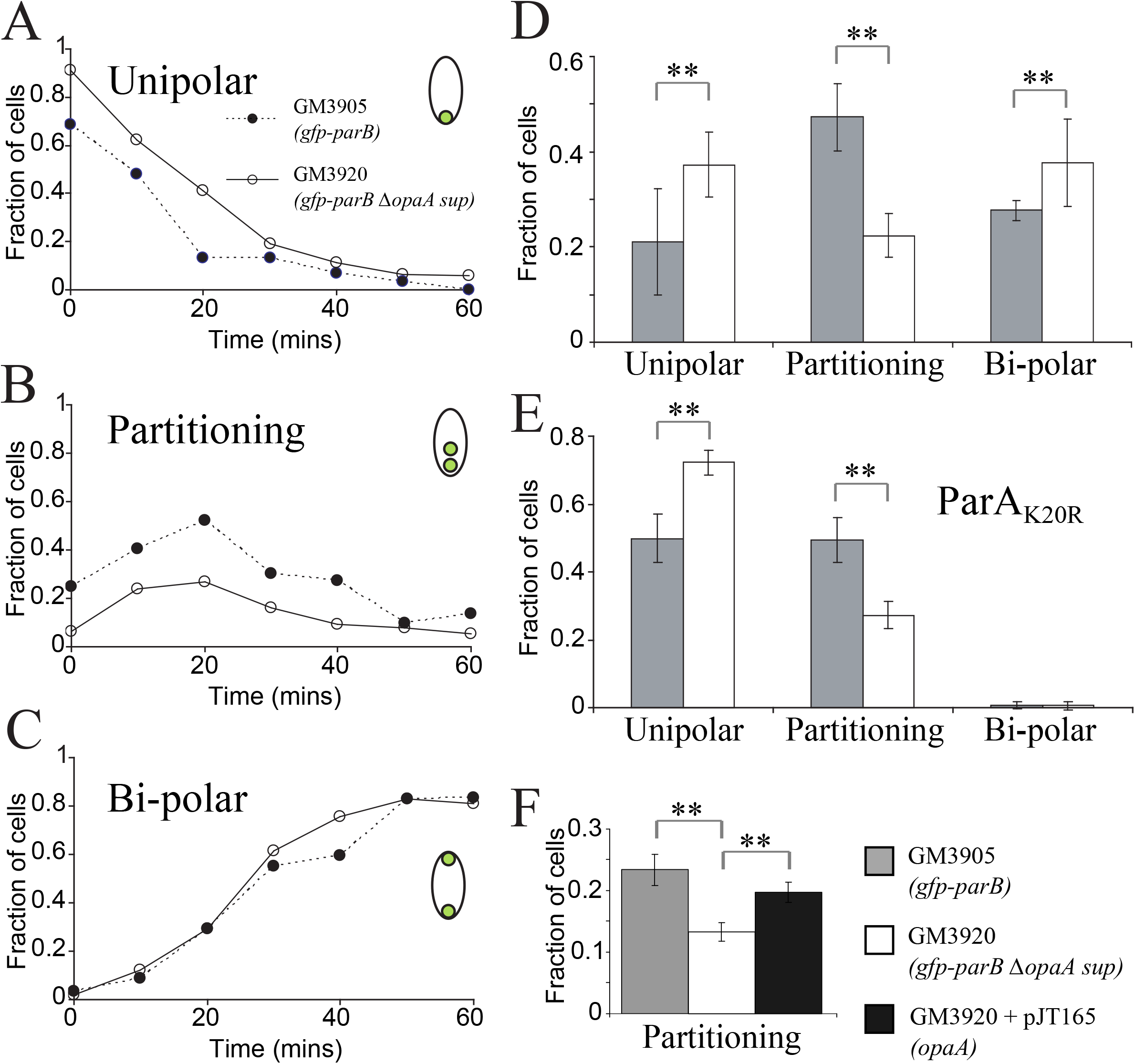
Null *opaA* cells are specifically defective in the early phase of chromosome partitioning. A-C) Quantification of cells at each stage of chromosome partitioning in synchronous cultures of WT cells carrying a chromosomal *gfp-parB* (GM3905; filled circles, dotted lines) or in a null *opaA* suppressor strain with the same chromosomal *gfp-parB* fusion (GM3920; empty circles, solid lines). Graphs show the changing number of cells that had a single GFP-ParB focus (A), a partially partitioned pair of GFP-ParB foci (B) or had completed partitioning (a pair of polar GFP-ParB foci; C) over the first 60 minutes of synchronous growth. n>100 cells for each time point. D) Quantification of cells at each stage of chromosome partitioning depicted in (A-C) at the t=20 minute time-point of three independent synchronous cultures of GM3905 (grey bars) or GM3920 (white bars). E) Quantification of cells at each stage of chromosome partitioning depicted in (A-C) at the t=20 minute time point of synchronous cultures of GM3905 (grey bars) and GM3920 (white bars) carrying a chromosomal *Pvan-parAK20R*. ParAK20R expression in these strains was induced two hours prior to synchronization and blocks ParA-mediated chromosome segregation. F) Quantification of cells in the process of partitioning chromosomes at t=25 minutes in GM3905 (grey bar), GM3920 (white bar) or GM3920 + pJT165 (carrying *opaA* under the native promoter; black bar) (n>300 for each condition). For D-F, n>300 cells for each condition. Significant differences (p<0.01, z-test) are indicated by **.

### Genetic analysis implicates opaA in early phase partitioning without ParA activity

The late phase of chromosome portioning requires ParA activity while the early phase does not [13]. To confirm that the defect in *parS*/GFP-ParB separation of GM3920 occurs during the early ParA-independent phase of chromosome segregation, we blocked ParA-mediated chromosome partitioning by expressing the dominant negative allele ParAK20R in synchronized cultures prior to the microscopic analysis at the 20 minute time point. As expected [13], expression of ParAK20R abolished bi-polar localization of GFP-ParB foci (Fig 5E) that requires ParA. However, the deficit of partially partitioned GFP-ParB foci remained in GM3920 when ParA was blocked, thereby conclusively placing the defect causing this deficit in the early ParA-independent phase of chromosome partitioning.

Because we examined chromosome partitioning in a suppressor strain rather than in a simple 317 *opaA* strain, we sought to confirm that the early partitioning deficit in GM3920 results from the lack of OpaA rather than from a suppressor mutation. We therefore transformed GM3920 with pJT165, carrying δ*opaA* under its native promoter to complement the *opaA* deletion, and we assayed chromosome partitioning as in Fig 5DE. pJT165 ameliorated the partitioning deficit of GM3920 (Fig 5F), further showing that OpaA directly aids the early phase of chromosome partitioning.

### Time-lapse microscopy implicates OpaA during earliest parS/centromere separation

To more closely investigate the deficit of double *parS/*GFP-ParB loci in δ*opaA* strain GM3920, we performed time-lapse microscopy on single cells of synchronized cultures. After 10 minutes of synchronous growth in PYE liquid, cells were transferred to nutrient PYE/agar pads on microscope slides for time-lapse analysis. Over the first 15 minutes of growth on the PYE/agar pads, fewer GM3920 cells started chromosome partitioning compared to control GM3905 cells (measured “start” events in Fig 6A). While implying a defective early separation, we judged these fewer “start” events as not statistically significant (P>0.05, z-test). However, statistically significant events, specifically fewer bi-polar “end” events (GM3920 cells that completed partitioning) and fewer “start and end” events (cells that both initiated and completed partitioning) were seen within the 15 minute period (Fig 6A).

**Fig 6.**
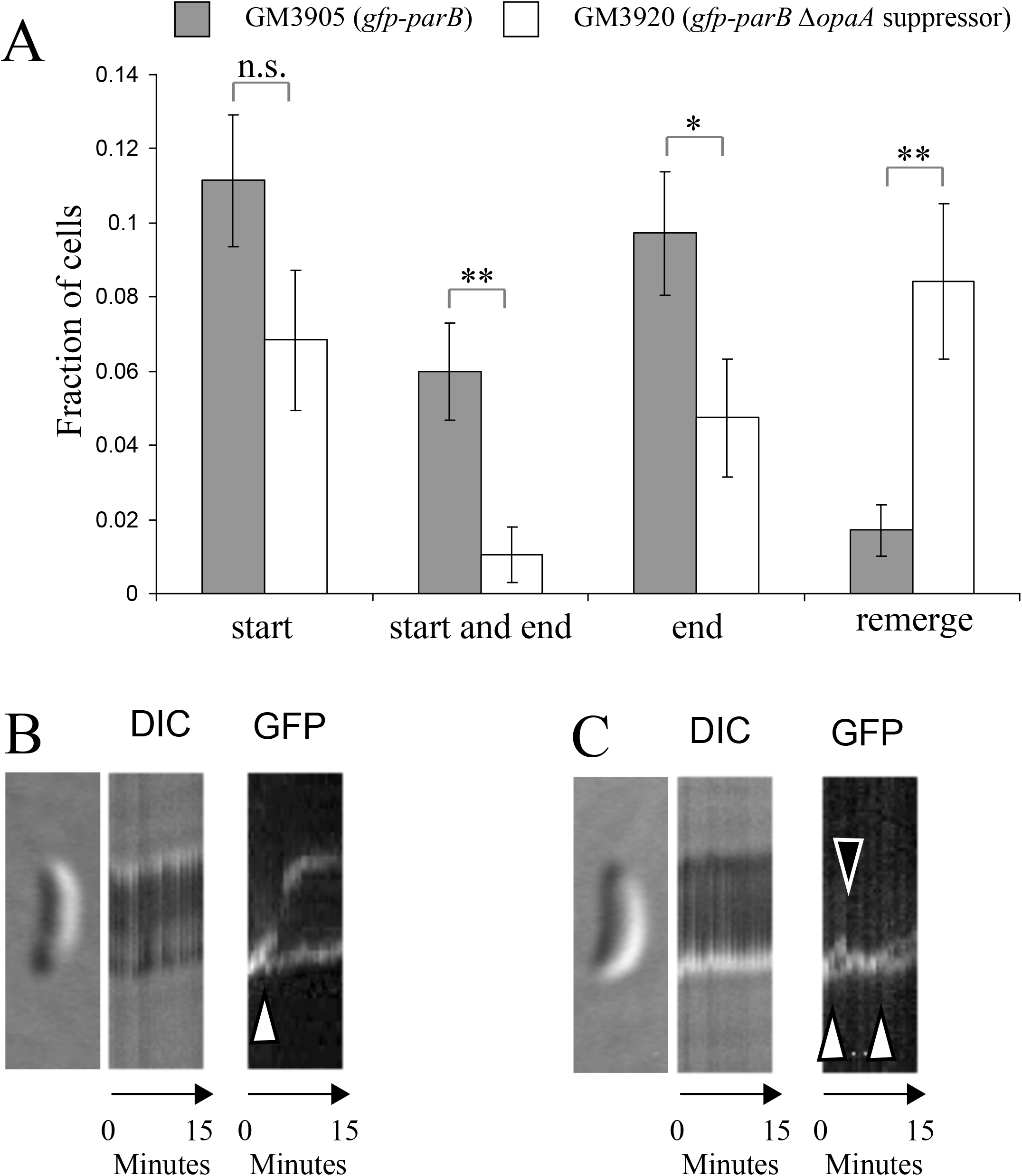
OpaA aids the early phase of chromosome partitioning. A) Quantification of the partitioning state of GM3905 (grey bars) and GM3920 cells (white bars) over a 15 minute time-course assayed by time-lapse microscopy. Cells were quantified according to whether they initiated partitioning (“start”; one polar GFP focus becomes two), completed partitioning (“end”; one polar and one mid-cell GFP focus become two bi-polar foci), both initiated and completed partitioning (“start and end”; one polar GFP focus becomes two bi-polar foci) or underwent a “remerge” event (two foci combined to show one focus) over the 15 minutes of time-lapse microscopy. Error bars indicate the estimated standard deviation of a Poisson distribution (n=350 and 190 for GM3905 and GM3920 respectively). Significant differences have probabilities (z-test) of p<0.05 and p<0.01, indicated by * and ** respectively. No significant difference (n.s.) p>0.05. B and C) Single-cell kymographs created from sliced and aligned time-lapse images, taken every minute over 15 minutes. Examples of GM3905 (B) and of GM3920 (C) cells undergoing a normal partitioning (B) and of a failed partitioning (C) of GFP-ParB foci. To the left of the DIC kymograph, the first whole cell image is indicated to show pole positions (note some drift in cell position during the experiment). White arrowheads in the GFP kymographs indicate separation events where two GFP-ParB foci became resolvable. The black arrowhead in the GFP kymograph in (C) indicates a “remerge” event where two GFP-foci fused into one foci. These images therefore illustrate the “start and end” (B) and “remerge” (C) events quantified in (A).

Most importantly, we also saw a significantly higher proportion of GM3920 cells where two GFP-ParB foci first separated but then “remerged” into one focus (Fig 6A, with a specific example in 6C). We interpret a “remerge” event as a failure to maintain the first separation step. This step coincides with what is conceptually a symmetry-splitting decision of the *par* system to keep one *parS* locus at the *St* pole and to send the other *parS* locus to the emerging *Sw* pole. These observations probably account for the deficit of separated GFP-ParB foci in GM3920 that we scored as “partitioning” foci in Fig 5. To the best of our knowledge, such *parS* remerging has not been reported before, probably because it is rare in WT cells (Fig 6A) and it only becomes apparent in our *opaA*-null cells when the early partitioning phase is closely examined.

## Discussion

Despite much research, bacterial chromosome replication and partitioning remain elusive processes. The many annotated genes with unknown functions argue that many DNA-binding proteins have not yet been found [14]. Here we found and characterized the novel DNA-binding protein, OpaA. We purified OpaA based on its binding to the most conserved and functionally important DNA inside the *C. crescentus* origin of chromosome replication (*Cori*) and we proposed that OpaA regulates chromosome replication, because increased OpaA binding to *Cori*UP mutant DNA *in vitro* correlates with increased autonomous replication of *Cori*UP mutant plasmids *in vivo* (Fig 1). Subsequent experiments support this chromosome replication hypothesis. For example, our genetic shut-off experiments show that lack of OpaA selectively prevents the initiation of chromosome replication *in vivo* (Fig 2). We scrutinized the consequences of *opaA* shut-off over the short-term 90 min period of the *C. crescentus* cell cycle. These are technically *in vivo* protein depletion experiments and since OpaA is a stable protein, we actually observed the consequences of reduced OpaA protein levels. Under these conditions and as in WT cells, swarmer (*Sw*) cells differentiated into stalked cells (*St*) which then proceeded with the asymmetric cell division. The CtrA master regulator marks the *Sw* cells and its disappearance marks the development of *St* cells while its reappearance (new synthesis) marks the onset of asymmetric cell division [15, 16]. Therefore, it is very significant that CtrA degradation kinetics are not changed by OpaA while the initiation of chromosome replication is severely retarded in the *opaA* shut-off and arrested in most individual cells (Fig 2BC, compare Pxyl::*opaA* strain GM3817 xylose versus glucose). Therefore, lack of OpaA selectively inhibits chromosome replication and we also presented two supplementary experiments (S4AB Fig) demonstrating that OpaA acts directly at *Cori*. Interestingly, OpaA protein levels increase during the cell cycle, and compared to *Sw* cells OpaA concentrations are at least 2-fold higher in dividing *St* cells (S9 Fig). Therefore, bulk OpaA protein concentrations are asymmetrically distributed by cell division and *Sw* cells receive lower concentrations of OpaA than *St* cells. However, OpaA protein concentrations alone do not determine DNA-binding. When OpaA concentrations are low, substantially more OpaA binds to the *parABS* DNA of synchronized *Sw* cells (Fig 3B). Also, OpaA binds to *Cori* with very specific spatial and temporal patterns (Fig 3B, compare *Cori* L, M and R patterns; Fig 1A). *Cori* is a platform for many protein interactions. Therefore, these OpaA binding patterns presumably reflect the replication fork assembly and regulatory interactions that occur during synchronized 380
chromosome replication.

How might OpaA regulate chromosome replication? *Cori* contains binding sites for DnaA and CtrA (Fig 1A) and previous studies have interpreted *Cori* regulation as an antagonism between positive DnaA activity and negative CtrA activity [7, 17, 18]. For example, high CtrA activity in *Sw* cells represses chromosome replication. Preliminary *in vitro* binding experiments with purified proteins show that OpaA protein preferentially displaces CtrA and not DnaA from *Cori* (data not shown). Yet antagonistic interactions with CtrA alone are not sufficient to explain the arrest of chromosome replication during OpaA depletion, as CtrA is still readily cleared in these cells (Fig 2D). Therefore OpaA likely modulates the binding of DnaA and probably additional regulators to *Cori*. The dynamic distribution of OpaA on *Cori* probably reflects complex interactions with additional regulators of replication (Fig 3B). In summary, all of our data are consistent with OpaA acting to promote chromosome replication.

How is OpaA involved in chromosome partitioning? We used ChIP-seq to identify the global distribution of OpaA binding and we found that the top ranked OpaA binding peaks were around the chromosomal partitioning locus (*par*; Fig 3; Table 1). We therefore addressed how chromosome partitioning could be affected by OpaA. In spite of several decades of research, the mechanisms by which bacterial chromosomes are partitioned remain incompletely understood. In many organisms, including *C. crescentus*, a *parABS* partitioning system, originally found on stable plasmids, is responsible for driving daughter chromosomes to opposite cell poles. While it has been shown that this system consists of a centromere-like *parS* locus that binds protein ParB, and an ATPase ParA that supplies the energy for *par*S-ParB translocation, the mechanism(s) of translocation remain(s) unclear.

Moreover, C*. crescentus* chromosome partitioning is bi-phasic, with an initial slow separation of *parS*-ParB complexes [13] followed by a rapid movement of one *parS*-ParB complex to the new cell pole [19]. Only the rapid second phase of partitioning requires ParA. Blocking ParA activity by expressing the dominant negative allele ParAK20R abolishes final *parS* placement at the distant cell pole [13, 19]. Nevertheless, these ParAK20R cells still undergo the initial slow and partial separation of *parS*-ParB. Our microscopy studies showed that significantly fewer cells separate the *parS*-ParB complexes in the *opaA* suppressor strain and our experiments placed the separation defect in the initial ParA-independent phase of chromosome partitioning. Additionally, time lapse microscopy showed that fewer δ*opaA* suppressor cells separate their *parS*-ParB complexes, and that a greater proportion of them remerge *parS-*ParB. Therefore, the initial separation of the *par-Cori* region is not maintained and OpaA may supply or help direct a driving force for this separation.

In this regard, the dynamic cellular mCherry-OpaA distributions (Fig 4) also argue that OpaA acts during the initial slow separation phase. Interestingly, the strongest mCherry-fluorescence stops next to the *par*S-ParB-GFP focus in the t=0 *Sw* cells (Fig 4). Subsequently, two *par*S-ParB-GFP foci form and one focus follows the mCherry-fluorescence as it recedes away from the cell pole still occupied by the remaining focus (t=15, t=30 cells). These mCherry-OpaA movements are independent of ParA (data not shown) and they suggest a separate quasi-mitotic process that makes many fluid contacts with the chromosome. At later times, the *par*S-ParB-GFP focus stops following and instead moves through the main mCherry-OpaA fluorescence on its way to the opposite cell pole. We suggest that this movement into the mCherry zone reflects a transfer from the early/slow OpaA-system to the later/fast ParA-system.

Therefore, mCherry-OpaA fluorescence microscopy suggests that OpaA movements could be a motor for chromosome movement. The motor responsible for driving the early phase of chromosome segregation is unknown [13] and previous studies have suggested RNA polymerase, DNA replication and bulk polymer properties may all account for directed chromosome movements [20-22]. Analysis of suppressor mutants can help identify the mechanistic interactions of essential movement-directing proteins. We therefore performed whole genome sequencing analysis on four independently derived δ*opaA* suppressor strains to identify candidate lethality suppressing mutations (S2 Table; supplementary methods). We found three independent mutations in the 23S rRNA gene and one mutation in *rpoB* encoding the RNA polymerase β-subunit, thereby supporting an interaction(s) between transcription/translation and OpaA. In *C. crescentus* translating ribosomes are not free to diffuse [23], so translating ribosomes can push the mRNA and DNA-template. Given also the link we describe between RNA polymerase activity and OpaA DNA binding (Fig 3C), might OpaA somehow link transcription/translation forces to the chromosomal DNA at *parB* in order to assist early chromosome movements? To test if transcriptional and translational forces are required during early partitioning, we examined partitioning in cells where transcription or translation had been disrupted, as follows: To focus on early partitioning, we used a *gfp-parB* strain expressing ParAK20R to poison ParA mediated partitioning. Then to separate replication from partitioning, we allowed synchronized cells to progress to a point where most cells had initiated replication and observed the localization of *gfp-parB* foci after brief treatment with antibiotics that disrupt transcription and translation. We saw that both rifampicin (an inhibitor of RNA polymerase) and chloramphenicol (an inhibitor of translational elongation) caused an increased number of cells with a single GFP-ParB-*parS* focus, which are presumably “remerged” double foci, because these cells initiated replication (S10 Fig). Therefore blockage of transcription/translation causes a partitioning phenotype similar to that seen in the δ*opaA* suppressor (Figs 5 and 6). These observations are consistent with forces exerted by transcription/translation acting during the early phase of chromosome segregation. Our data therefore suggest a link between transcription/translation and the function of OpaA (suppressor mutations S2 Table; ChIP-seq data Fig 3C; microscopy data S10 Fig). To confirm that 23S rRNA mutations can suppress δ*opaA* lethality, we isolated 10 additional spontaneous suppressor mutants and when we sequenced their 23S rRNA genes using PCR, we found an additional 3 independent point mutations in the 23S rRNA (data not shown). There is also precedent for ribosome components regulating replication [24]. Therefore for future studies, it will be especially important to understand the link between the ribosome, chromosome replication and chromosome movements that is implied by these mutations.

Bacterial chromosome replication and partition overlap temporally and this regulatory link is also implied by chromosome organization, because origins of replication often lie close to *par* loci. For example, in *C. crescentus Cori* and *parS* are separated by only 10 kb and the 1st, 2nd and 3rd ranked OpaA-binding peaks all span this region (Table 1). These temporal and spatial linkages probably evolved to meet the special needs of bacteria. Theoretically, it should also be advantageous for both *par*S and *Cori* to share regulatory proteins. Interestingly, a recent study also implicated the replication initiator DnaA during chromosome partitioning in *C. crescentus* [25]. Therefore, replication and partitioning both utilize DnaA and OpaA in *C. crescentus*. Similarly, in *Vibrio cholerae* and *Bacillus subtillis*, ParA interacts with DnaA to control replication [26, 27]. Since chromosome replication precedes its partitioning, one might expect regulators to act sequentially, first at *Cori* and then at *parS*, but counter intuitively a recent study shows that as the activity of *C. crescentus* DnaA rises, it first acts at *parS* and then at *Cori*, because the threshold for DnaA action is lower at *parS* than at *Cori* [25]. In a like manner, we see peak binding of OpaA at *parB* immediately preceding the OpaA binding interactions at *Cori* (*Cori* M and R) during the initiation of replication (Fig 3B). Therefore, while earlier cell cycle binding of OpaA to *par*S at first appeared to be a paradoxical attribute of a replication regulator, it now agrees with an emerging view that both OpaA and DnaA must respond to cell cycle signals and by as yet unknown mechanisms they must prepare *par*S in anticipation of replication initiation at *Cori*.

While the exact mechanism remains to be fully explored, OpaA clearly acts during the critical period of the cell cycle when chromosome replication and partitioning are initiated. The early ParA-independent separation of *parS* loci is very important for asymmetric cell division, because this separation enables ParA, which moves from the distal pole, to select just one chromosome for transport. If two *parS*-ParB loci are in close proximity when they encounter ParA, they risk being co-transported and not segregated [13, 28, 29]. Therefore, while the ParA-system acts indiscriminately, it is the earlier OpaA-ultilizing system that discriminates between two alternative chromosomes. The initial separation of *parS* does not absolutely require OpaA, as it still occurs in the null *opaA* suppressor strain, but OpaA clearly aids in initiating and maintaining this separation (Figs 5 and 6). To our knowledge OpaA is the first specific protein identified with the early ParA-independent phase of chromosome partitioning. This is a key symmetry-splitting phase that channels the chromosomes into distinct *Sw*-pole versus *St*-pole programs (including alternative chromosome replication programs). Therefore, we propose that OpaA facilities *C. crescentus* asymmetry. This role may also explain why OpaA homologs are found exclusively in the α-proteobacteria [30] which typically divide asymmetrically and localize their origins of replication at the cell poles [31-33].

In summary, OpaA is a novel protein that facilitates at least two molecular “decision-making” systems during the one of the most critical parts of the *C. crescentus* cell cycle: The “decision” to initiate chromosome replication and the “decision” to partition just one chromosome to the newly developing (incipient *Sw*) cell pole. Understanding how OpaA interacts with their molecular components, e.g. through genetic-suppressor and biochemical interaction studies should provide new insights into both of these fundamental and universal processes.

## Materials and Methods

### Bacterial strains and plasmids

*C. crescentus* strains and plasmids used in this study are described in S3 and S4 Tables, respectively. Construction of new plasmids and *C. crescentus* strains is described in the supplementary material. *E. coli* strains were grown in LB media supplemented with ampicillin 514 (100 μg/ml) where noted. *C. crescentus* strains were grown in PYE or g/ml), M2G media as noted, with xylose (0.5% w/v), glucose (0.2% w/v), ampicillin (20 μg/ml chloramphenicol (1 μg/ml), spectinomycin (100 μg/ml) and streptomycin (2.5 μg/ml) added as described.

### Cell fractionation and protein purification

The initial cell fractionation carried out to purify OpaA from whole cell lysates as well as the purification buffer compositions are described in S1 Fig. Recombinant His-OpaA and GST-CtrA were purified as described in supplementary material.

### EMSA Reactions

EMSA reactions were performed with radio-labeled oligonucleotides as previously described [7] but in EMSA buffer consisting of 20 mM Tris-HCl pH 8, 100 mM KCl, 5 mM MgCl2, 1 mM CaCl2, 2 mM DTT, 50 μg ml-1 BSA. Briefly, radio-labeled annealed oligonucleotides were incubated with various protein fractions in the EMSA buffer on ice for 30 min before being loaded directly onto a 8% polyacrylamide gel in 1x TBE. The sequences of the oligonucleotides used for *Cori*WT and *Cori*UP are listed in S5 Table (“WT” and “UP” pairs respectively).

### Autonomous replication assay

Autonomous replication assays were preformed as previously described [7]. Briefly, plasmids carrying the reporter gene *gusA* and relying on *Cori* or *Cori*UP for their replication were transformed by electroporation into GM1609 or derivatives, and cells were plated onto selective media (PYE + ampicillin). After four days, colonies were washed off into PYE media, and assayed for GusA activity to report plasmid copy number.

### Immunoblotting

Antibodies were raised in rabbits against OpaA (at the McGill Animal Facility) and CtrA (at MediMabs, Montreal), or have previously been described [9]. All were used in Western blotting at a 1:10,000 dilution. Secondary anti-rabbit antibodies conjugated to horse radish peroxidase (Sigma) were also used at a 1:10,000 dilution. Immunoblots from SDS-PAGE onto PVDF membrane were visualized using Western Lightning reagents (Perkin-Elmer) and a VersaDoc system (BioRad).

### Chromatin ImmunoPrecipitation coupled to deep Sequencing (ChIP-Seq)

ChIP-seq was carried out essentially as previously described [34] using mid-log phase cells (O.D.660nm~0.5), cultivated in PYE or preincubated for 10 min with antibiotics (Rifampicin 30μg/ml; Novobiocin 100μg/ml). HiSeq 2000 runs of barcoded ChIP-Seq libraries yielded several million reads that were mapped to the *Caulobacter crescentus* NA1000 (NC_011916, circular form) and analyzed as described in supplementary methods. Briefly, the genome was subdivided into 1 bp (isolated regions) or 50 bp (full chromosome) probes, and for every probe we calculated the percentage of reads per probe as a function of the total number of reads. Analyzed data illustrated in Fig 3A using the Circos Software [35] are provided in S1 Dataset (50 bp resolution). Fig 3B focuses on the *par* and the *Cori* regions (4026155 to 1150 bp on the circular *Caulobacter crescentus* genome), analyzed datasets are provided in S2 Dataset (full chromosome at 50 bp resolution) and S3 Dataset (*par* and *Cori* regions at 1 bp resolution).

### Quantitative Chromatin Immunoprecipitation (qChIP)

Chromatin immunoprecipitation was performed as previously described but using 1:1000 anti-OpaA [34]. For mixed culture experiments, log phase cultures in PYE were treated with antibiotics (where described) for 10 minutes. qPCR was performed with oligonucleotide primers described in S5 Table (primer pairs listed as “q[gene name] fwd/rev”) using the FastStart Universal SYBR green qPCR kit (Roche) on a Rotorgene 6000 thermocycler (Corbett) or a CFX thermocycler (Bio-Rad).

### Flow cytometry

Flow cytometry was performed on synchronous cultures that had been isolated with a Percoll density gradient [9] using an out-growth protocol that allowed cells to finish any rounds of chromosome replication that had already initiated, as previously described [2]. Briefly, freshly isolated swarmer cells were transferred to PYE containing glucose or xylose and samples were taken from the synchronized culture at various times and transferred into media containing 60 μg/ml cephalexin for at least three hours. Ethanol was then added to 70% final concentration to fix cells, and samples washed once with 70% ethanol and were stored at 4 °C prior to analysis. Staining and analysis were performed as described [2].

### Microscopy

Fluorescence and DIC images were captured using a Zeiss Axiovert200M with a 100x objective at the McGill University Life Sciences Complex Advanced BioImaging Facility. For fluorescence microscopy experiments, samples were imaged on 1% agarose pads in water or PYE media. For all experiments comparing GFP-ParB localization in synchronized cells, run out flow cytometry samples (see above) were collected concurrently to confirm that replication initiation levels were comparable between strains. Determination of spot localization was performed manually on large numbers of cells (n-values are given in the respective figure legends).

## Acknowledgements

We also thank Drs. S. Sagan, C. Maurice and T. Rolain for constructive criticisms of this manuscript.

## Supporting Information

**S1 Text. Containing supplementary methods, tables, figure legends and references.**

**S1 Fig. Purification of a *Cori* binding protein.**

**S2 Fig. OpaA depletion causes cell death.**

**S3 Fig. Degradation of cell cycle marker protein McpA.**

**S4 Fig. Direct *in vivo* binding of OpaA to *Cori* affects replication.**

**S5 Fig. qChIP experiments supporting ChIP-seq data in Fig 2.**

**S6 Fig. Preliminary analysis of the DNA binding determinants in OpaA.**

**S7 Fig. ChIP-seq plots surrounding additional selected peaks of OpaA binding.**

**S8 Fig. A complete set of images, a subset of which are shown in Fig 7.**

**S9 Fig. Western blot showing OpaA levels over the cell.**

**S10 Fig. Quantification of the location of *parS*.**

**S11 Fig. A phylogenetic tree generated using Clustal Omega.**

**S1_Figs. Containing corresponding supplementary figures.**

## References

1. Bartosik AA, Jagura-Burdzy G. Bacterial chromosome segregation. Acta biochimica Polonica. 2005;52(1):1–34.

2. Bastedo DP, Marczynski GT. CtrA response regulator binding to the Caulobacter chromosome replication origin is required during nutrient and antibiotic stress as well as during cell cycle progression. Mol Microbiol. 2009;72(1):139–54.

3. Wolanski M, Donczew R, Zawilak-Pawlik A, Zakrzewska-Czerwinska J. oriC-encoded instructions for the initiation of bacterial chromosome replication. Frontiers in microbiology. 2014;5:735.

4. Marczynski GT, Rolain T, Taylor JA. Redefining bacterial origins of replication as centralized information processors. Front Microbiol. 2015;6:610.

5. Laub MT, Shapiro L, McAdams HH. Systems biology of Caulobacter. Annu Rev Genet. 2007;41:429–41.

6. Mohl DA, Easter J, Jr., Gober JW. The chromosome partitioning protein, ParB, is required for cytokinesis in Caulobacter crescentus. Mol Microbiol. 2001;42(3):741–55.

7. Taylor JA, Ouimet MC, Wargachuk R, Marczynski GT. The Caulobacter crescentus chromosome replication origin evolved two classes of weak DnaA binding sites. Mol Microbiol. 2011;82(2):312–26.

8. Christen B, Abeliuk E, Collier JM, Kalogeraki VS, Passarelli B, Coller JA, et al. The essential genome of a bacterium. Molecular systems biology. 2011;7:528.

9. Tsai JW, Alley MR. Proteolysis of the Caulobacter McpA chemoreceptor is cell cycle regulated by a ClpX-dependent pathway. J Bacteriol. 2001;183(17):5001–7.

10. Bailey TL, Williams N, Misleh C, Li WW. MEME: discovering and analyzing DNA and protein sequence motifs. Nucleic Acids Res. 2006;34(Web Server issue):W369–73.

11. Fumeaux C, Radhakrishnan SK, Ardissone S, Theraulaz L, Frandi A, Martins D, et al. Cell cycle transition from S-phase to G1 in Caulobacter is mediated by ancestral virulence regulators. Nature communications. 2014;5:4081.

12. Thanbichler M, Shapiro L. MipZ, a spatial regulator coordinating chromosome segregation with cell division in Caulobacter. Cell. 2006;126(1):147–62.

13. Shebelut CW, Guberman JM, van Teeffelen, S. Yakhnina AA, Gitai Z. Caulobacter chromosome segregation is an ordered multistep process. Proc Natl Acad Sci U S A. 2010;107(32):14194–8.

14. Rigden DJ. Ab initio modeling led annotation suggests nucleic acid binding function for many DUFs. OMICS. 2011;15(7-8):431–8.

15. Quon KC, Marczynski GT, Shapiro L. Cell cycle control by an essential bacterial two-component signal transduction protein. Cell. 1996;84(1):83–93.

16. Domian IJ, Quon KC, Shapiro L. Cell type-specific phosphorylation and proteolysis of a transcriptional regulator controls the G1-to-S transition in a bacterial cell cycle. Cell. 1997;90(3):415–24.

17. Quon KC, Yang B, Domian IJ, Shapiro L, Marczynski GT. Negative control of bacterial DNA replication by a cell cycle regulatory protein that binds at the chromosome origin. Proc Natl Acad Sci U S A. 1998;95(1):120–5.

18. Jonas K, Chen YE, Laub MT. Modularity of the bacterial cell cycle enables independent spatial and temporal control of DNA replication. Current biology: CB. 2011;21(13):1092–101.

19. Toro E, Hong SH, McAdams HH, Shapiro L. Caulobacter requires a dedicated mechanism to initiate chromosome segregation. Proc Natl Acad Sci U S A. 2008;105(40):15435–40.

20. Lemon KP, Grossman AD. Movement of replicating DNA through a stationary replisome. Mol Cell. 2000;6(6):1321–30.

21. Jun S, Mulder B. Entropy-driven spatial organization of highly confined polymers: lessons for the bacterial chromosome. Proc Natl Acad Sci U S A. 2006;103(33):12388–93.

22. Dworkin J, Losick R. Does RNA polymerase help drive chromosome segregation in bacteria? Proc Natl Acad Sci U S A. 2002;99(22):14089–94.

23. Montero Llopis P, Sliusarenko O, Heinritz J, Jacobs-Wagner C. In vivo biochemistry in bacterial cells using FRAP: insight into the translation cycle. Biophysical journal. 2012;103(9):1848–59.

24. Chodavarapu S, Felczak MM, Kaguni JM. Two forms of ribosomal protein L2 of Escherichia coli that inhibit DnaA in DNA replication. Nucleic Acids Res. 2011;39(10):4180–91.

25. Mera PE, Kalogeraki VS, Shapiro L. Replication initiator DnaA binds at the Caulobacter centromere and enables chromosome segregation. Proc Natl Acad Sci U S A. 2014;111(45):16100–5.

26. Kadoya R, Baek JH, Sarker A, Chattoraj DK. Participation of chromosome segregation protein ParAI of Vibrio cholerae in chromosome replication. J Bacteriol. 2011;193(7):1504–14.

27. Murray H, Errington J. Dynamic control of the DNA replication initiation protein DnaA by Soj/ParA. Cell. 2008;135(1):74–84.

28. Hwang LC, Vecchiarelli AG, Han YW, Mizuuchi M, Harada Y, Funnell BE, et al. ParA-mediated plasmid partition driven by protein pattern self-organization. EMBO J. 2013;32(9):1238–49.

29. Ptacin JL, Gahlmann A, Bowman GR, Perez AM, von Diezmann AR, Eckart MR, et al. Bacterial scaffold directs pole-specific centromere segregation. Proc Natl Acad Sci U S A. 2014;111(19):E2046–55.

30. Gupta RS, Mok A. Phylogenomics and signature proteins for the alpha proteobacteria and its main groups. BMC Microbiol. 2007;7:106.

31. Jensen RB, Wang SC, Shapiro L. Dynamic localization of proteins and DNA during a bacterial cell cycle. Nature reviews Molecular cell biology. 2002;3(3):167–76.

32. Deghelt M, Mullier C, Sternon JF, Francis N, Laloux G, Dotreppe D, et al. G1-arrested newborn cells are the predominant infectious form of the pathogen Brucella abortus. Nature communications. 2014;5:4366.

33. Kahng LS, Shapiro L. Polar localization of replicon origins in the multipartite genomes of Agrobacterium tumefaciens and Sinorhizobium meliloti. J Bacteriol. 2003;185(11):3384–91.

34. Quisel JD, Lin DC, Grossman AD. Control of development by altered localization of a transcription factor in B. subtilis. Mol Cell. 1999;4(5):665–72.

35. Krzywinski M, Schein J, Birol I, Connors J, Gascoyne R, Horsman D, et al. Circos: an information aesthetic for comparative genomics. Genome Res. 2009;19(9):1639–45.

